# Kasugamycin is a novel chitinase 1 inhibitor with strong antifibrotic effects on pulmonary fibrosis

**DOI:** 10.1101/2021.02.25.432796

**Authors:** Jae-Hyun Lee, Chang-Min Lee, Joyce H. Lee, Mun-Ock Kim, Jin Wook Park, Suchitra Kamle, Bedia Akosman, Erica L. Herzog, Xue Yan Peng, Jack A. Elias, Chun Geun Lee

## Abstract

**Rationale:** Pulmonary fibrosis is a devastating lung disease with few therapeutic options. Chitinase 1 (CHIT1), an 18 glycosyl hydrolase family member, contributes to the pathogenesis of pulmonary fibrosis through regulation of Transforming Growth Factor (TGF)-β signaling and effector function. Therefore, CHIT1 is a potential therapeutic target of pulmonary fibrosis.

**Objectives:** This study aimed to identify and characterize a druggable CHIT1 inhibitor with strong antifibrotic activity and minimal toxicity for therapeutic application to pulmonary fibrosis.

**Methods:** Extensive screening of small molecule libraries identified the aminoglycoside antibiotic Kasugamycin as a potent CHIT1 inhibitor.

**Measurements and Main Results:** Elevated levels of CHIT1 were detected in the lungs of patients with pulmonary fibrosis. In vivo bleomycin- and TGF-β-stimulated murine models of pulmonary fibrosis, Kasugamycin showed impressive anti-fibrotic effects in both preventive and therapeutic conditions. In vitro studies also demonstrated that Kasugamycin inhibits fibrotic macrophage activation, fibroblast proliferation and myofibroblast transformation. Null mutation of transforming growth factor beta associated protein 1 (TGFBRAP1), a recently identified CHIT1 interacting signaling molecule, phenocopied antifibrotic effects of Kasugamycin in in vivo lungs and in vitro fibroblasts responses. Kasugamycin inhibits physical association between CHIT1 and TGFBRAP1, suggesting that antifibrotic effect of Kasugamycin is mediated through regulation of TGFBRAP1, at least in part.

**Conclusions:** These studies demonstrate that Kasugamycin is a novel CHIT1 inhibitor with strong antifibrotic effect that can be further developed as an effective and safe therapeutic drug for pulmonary fibrosis.

## INTRODUCTION

The 18 Glycosyl hydrolase (18GH) gene family includes a number of chitinase and chitinase-like proteins (1). Among these, chitinase 1 (CHIT1) and acidic mammalian chitinase (AMCase) are true chitinases that bind and cleave chitin. Chitinases are found in a variety of species ranging from lower life forms (archea, prokaryotes, eukaryotes) to humans, and they are known to play a critical role in the life cycle of chitin-containing organisms including insects. CHIT1 is the major chitinase noted in mammals including humans, however, the exact biological function of CHIT1 in mammals is not clear since chitin and chitin synthase do not exist in mammals and higher life forms do not use chitin as a nutrient (2, 3).

Recently, we reported that CHIT1 plays a significant role in the pathogenesis of interstitial lung disease associated with scleroderma (SSc-ILD) and idiopathic pulmonary fibrosis (IPF) (4, 5). The expression of CHIT1 is induced in the lungs of mice challenged with bleomycin or transforming growth factor-beta (TGF-β) overexpression and the increased lung fibrosis was abrogated in the mice with null mutations of CHIT1 (*Chit1*^-/-^) but further enhanced in the mice with CHIT1 overexpression (4). *In vitro* studies using murine alveolar macrophages and normal human lung fibroblasts demonstrated that CHIT1 is required and sufficient for profibrotic macrophage activation, fibroblast proliferation and myofibroblast transformation (5). These studies strongly support an essential role of CHIT1 in the pathogenesis of pulmonary fibrosis and CHIT1 could be a reasonable therapeutic target of pulmonary fibrosis.

In an effort to search for therapeutic antifibrotic drug(s) for intervention of pulmonary fibrosis based on anti-CHIT1 activity, we employed high throughput small molecule screening that includes multiple libraries of small molecules with druggable potential. After screening of >7500 small molecules, we identified that aminoglycoside antibiotic Kasugamycin (KSM) is a specific CHIT1 inhibitor with strong anti-fibrotic effects on bleomycin- and TGF-β-stimulated pulmonary fibrosis. *In vitro* studies demonstrated that KSM inhibits profibrotic macrophage activation, fibroblast proliferation and myofibroblast transformation. Co-immunoprecipitation and immunoblot assays further revealed that KSM inhibits the physical association between CHIT1 and TGFBRAP1, a CHIT1-interacting molecule known to enhance TGF-β signaling by acting as a chaperone of SMAD4 (5, 6). Taken together, these studies demonstrate that KSM is a novel CHIT1 inhibitor with strong antifibrotic activity that significantly inhibits the development and or progression of pulmonary fibrosis.

## RESULTS

### Dysregulated expression of CHIT1 in the lungs of IPF patients

To evaluate CHIT1 expression in the lungs of patients with idiopathic pulmonary fibrosis (IPF), immunohistochemistry (IHC) was performed on the archived lung biopsy specimens obtained from the Yale Lung Repository using anti-human polyclonal CHIT1 antibody as previously described by our laboratory (4). Histologically normal lungs were obtained from the surgical margin of lung biopsies for nodules that were later found to be benign. The number of CHIT1 positive cells (brown stain) were significantly higher in the lungs of IPF patients compared to normal lungs and macrophages and epithelial-like cells are the major cells expressing CHIT1 (Figure 1, A and B). Modestly increased expression of CHIT1 was also noted in other interstitial parenchymal cells in the lungs of IPF patients. Interestingly, double immunohistochemical evaluations identified that the CHIT1(+) expressing cells are only partially overlaps with F4/80 macrophages marker (Figure 1C, upper panel) but not with SPC or CC10 (Figure 1C and data not shown), suggest that macrophages with potentially altered surface marker expression are the major CHIT1 expressing cells in the lungs of IPF patients. These studies demonstrate that the expression of CHIT1 is dysregulated in the lungs of IPF patients.

**Figure 1.**
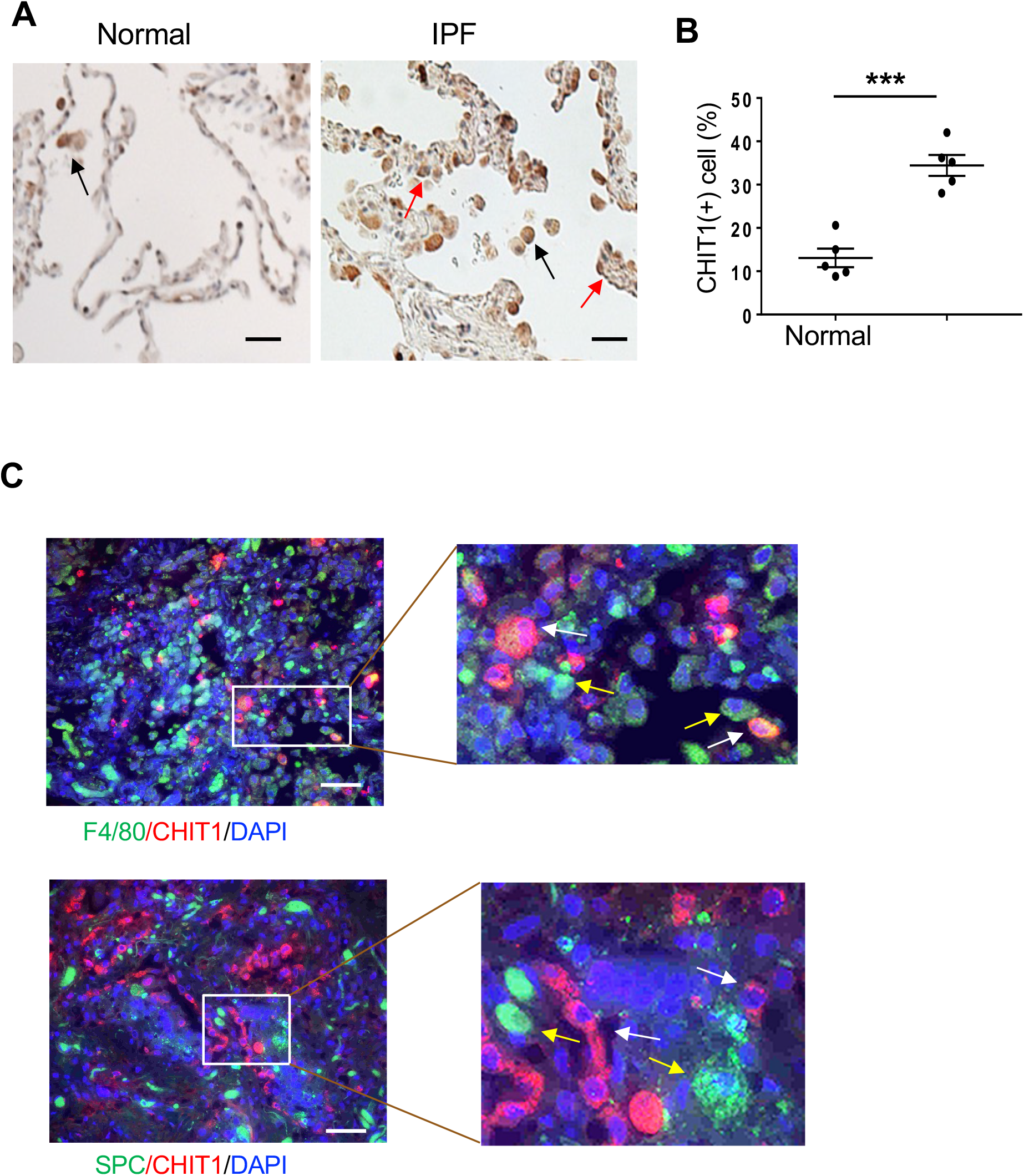
Dysregulated expression of CHIT1 in the lung of IPF patients. (A) Representative immunohistochemical staining of CHIT1 in the lung of normal and IPF patients. Arrows in panel A highlight CHIT1 stain (+) cells (black arrows, macrophages; red arrows, epithelial cells). (B) Quantitation of CHIT1 positive (+) cells in the lung sections from normal and IPF patient per high-powered field (20x of original magnification). (C) Double label immunohistochemistry (IHC) comparing localization of CHIT1, F4/80 and SPC expression in IPF lungs. White arrows indicate the CHIIT1 stain (+) cells and yellow arrows indicate the F4/80(+) macrophages (upper panel) and SPC (+) alveolar epithelial cells (lower panel). Each value in panels B is from a different individual and the mean±SEM are illustrated. \*\*\**P*<0.001 (Mann-Whitney U test). Scale bar in panel A, 50μm.

### Identification of Kasugamycin as a specific CHIT1 inhibitor

To identify druggable CHIT1 inhibitor, we used high throughput screening of small molecule libraries through the Yale Center for Molecular Discovery. This screening identified 42 compounds that significantly suppress catalytic activity of CHIT1 (>50% inhibition of CHIT1 activity compared to 200μM pentoxifylline, a known chitinase inhibitor (7)) (Table E1). Notably, these agents include several anti-cancer drugs and specific kinase inhibitors that are already reported to have certain antifibrotic effects, such as histone deacetylase (HDAC) inhibitor (CUDC-907) or tyrosine kinase inhibitor (Bosutinib) (8, 9). After careful chemistry review of each compound for potential toxicity (adverse effects) and validation kinetic studies (dose-dependent CHIT1 inhibition assay), KSM was selected as the most promising drug candidate based on strong anti-CHIT1 activity with no or low levels of proven animal toxicity (10, 11) (Figure 2A). KSM was first developed as an antibiotic humans in 1965 (12), since then *in vivo* pharmacokinetics and toxicology studies on KSM have been well established through human clinical and extensive animal toxicity studies (12–15). These studies demonstrated remarkable low toxicity of KSM in humans and in rodent toxicity studies in short and long term use. The anti-CHIT1 activity of KSM is specific among aminoglycoside antibiotics, since other members of aminoglycosides did not show anti-CHIT1 activity (Figure 2B).

**Figure 2.**
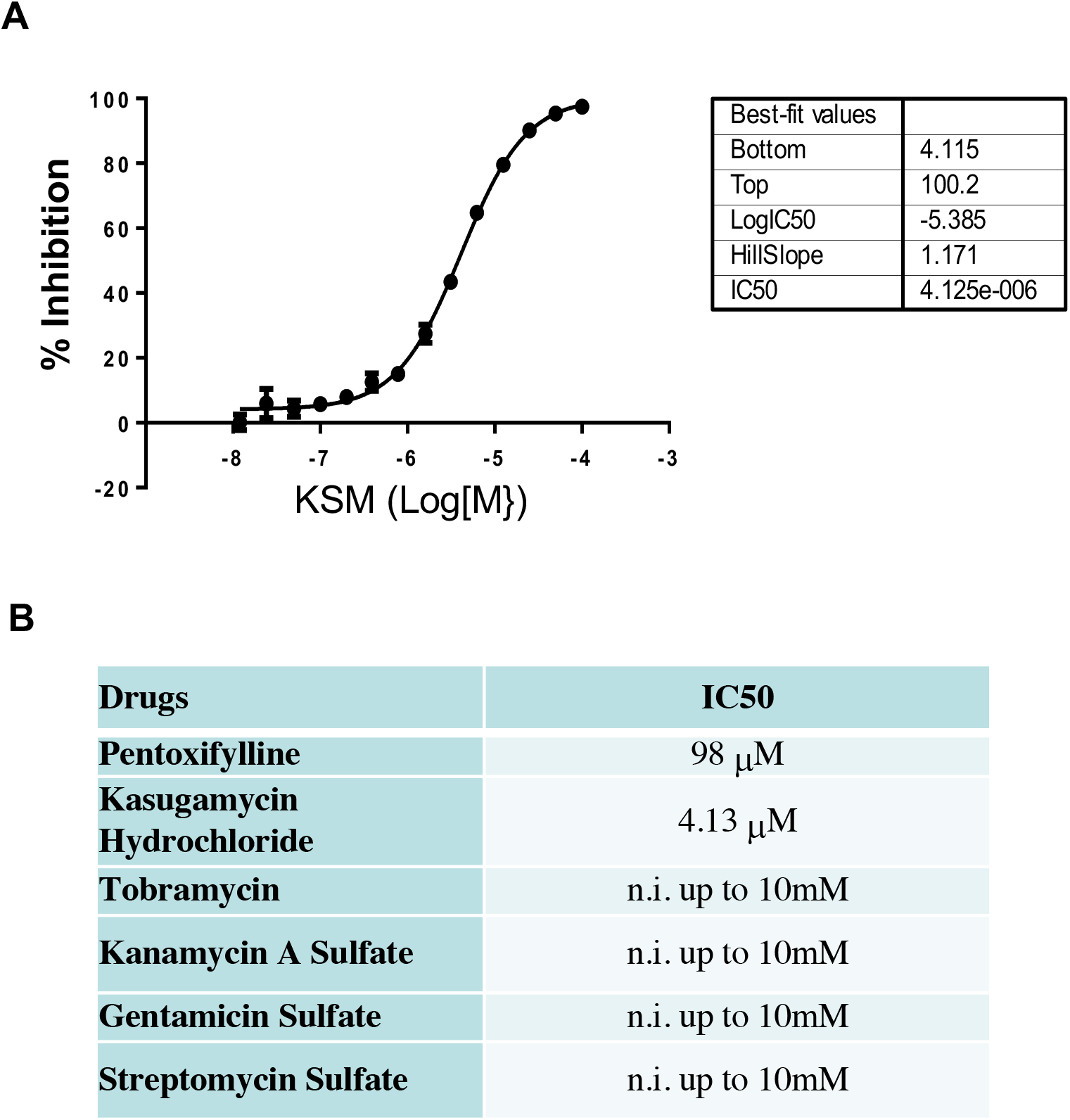
Effect of Kasugamycin and selected aminoglycosides on the CHIT1 activity. (A) Dose response curve of Kasugamycin (KSM) inhibition on the CHIT1 activity. (B) The effects of pentoxifylline and selected aminoglycosides on CHIT1 activity detected in small molecule library screening. Pentoxiphylline, a known CHIT1 inhibitor, was used as a positive reference for library screening. IC50, half maximal inhibitory concentration; n.i., no inhibition.

### KSM inhibits bleomycin-stimulated pulmonary fibrosis *in vivo*

Studies were initiated to test whether KSM has an antifibrotic effect in bleomycin-stimulated animal model of pulmonary fibrosis. In this study, preventive and therapeutic effects of KSM were tested by adjusting the timing of KSM delivery during or after bleomycin challenge. To ascertain preventive effects of KSM, different doses of KSM (12.5mg/kg to 100mg/kg, i.p., every other day) were administered to the mice during bleomycin challenge as illustrated in Figure 3A. On day 30 after first challenge of bleomycin, mice were sacrificed and the total collagen accumulation in the lung was evaluated by Sircol collagen assay and Mallory’s trichrome staining (Figure 3, B and C). These studies demonstrated that bleomycin-induced collagen accumulation in the lung was impressively decreased in the mice treated with KSM compared to vehicle controls in a dose dependent fashion (Figure 3, B and C). In accord with histologic changes, KSM inhibits bleomycin-stimulated expression of extracellular matrix (ECM) proteins including collagens (Col1a1 and Col3a1) and fibronectin (Figure 3D). Next, studies were undertaken to see whether KSM has a therapeutic benefit by evaluating the lungs of the mice treated with KSM or vehicle after establishment of fibrosis in the lung. In this evaluation, the mice were challenged with bleomycin for 6 consecutive days according to the previously reported protocol with minor modification (16, 17), then KSM or vehicle were delivered (every other day, 50mg/kg of KSM or vehicle, i.p.) to the mice from12 days after the first challenge of the bleomycin (Figure 4A). On day 25, the mice were sacrificed and fibrotic tissue responses in the lung were evaluated. In this therapeutic model of pulmonary fibrosis, bleomycin-stimulated collagen accumulation and the expression of ECM-associated genes in the lungs were abrogated in the mice treated with KSM compared to vehicle challenged ones (Figure 4, B and C). Accordingly, the bleomycin-stimulated ECM gene expression in the lung was also significantly reduced by KSM treatment (Figure 4D). These studies demonstrated that KSM has a strong anti-fibrotic activity against bleomycin-stimulated pulmonary fibrosis both in preventive and therapeutic conditions.

**Figure 3.**
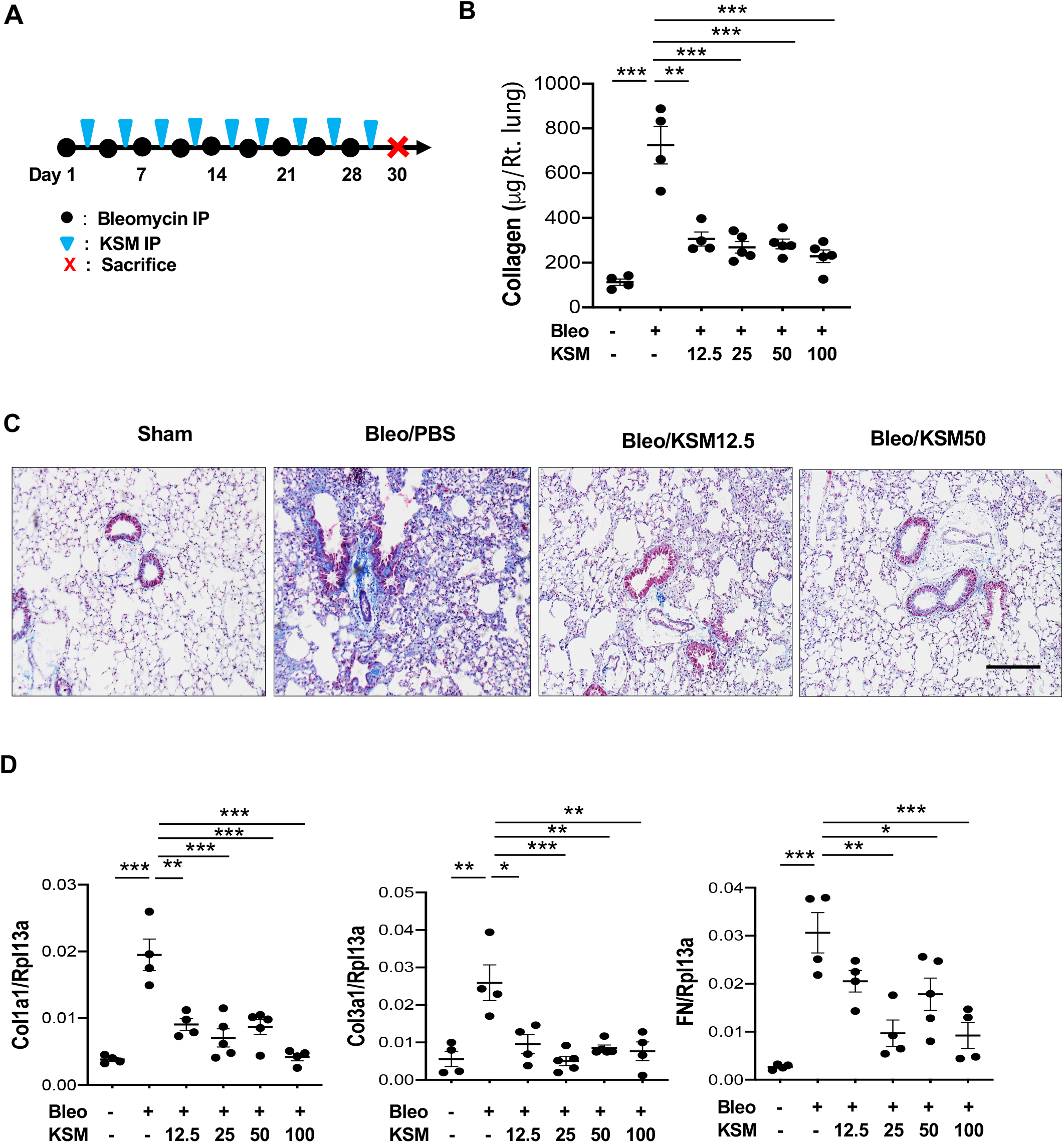
Kasugamycin inhibits bleomycin-stimulated pulmonary fibrosis. 6**-**8 weeks old male mice were challenged with bleomycin (Bleo; i.p, 0.25U/mouse) together with different doses of Kasugamycin (KSM) (12.5 to 100 mg/kg/mouse). (A) Schematic illustration of the protocol employed in this study. (B) Sircol collagen quantitation in the lungs. (C) Representative histology with Mallory’s trichrome staining. (D) Real-time qPCR evaluation of collagen I (Col1a1), III (Col3a1) and fibronectin (FN) expression in the lung. Rpl13a was used as an internal control. Each value in panels B and D is from a different animal and the mean±SEM are illustrated. \**P*<0.05, \*\**P*<0.01, \*\*\**P*<0.001 (One-Way ANOVA with multiple comparisons), Scale bars in panel C=50μm.

**Figure 4.**
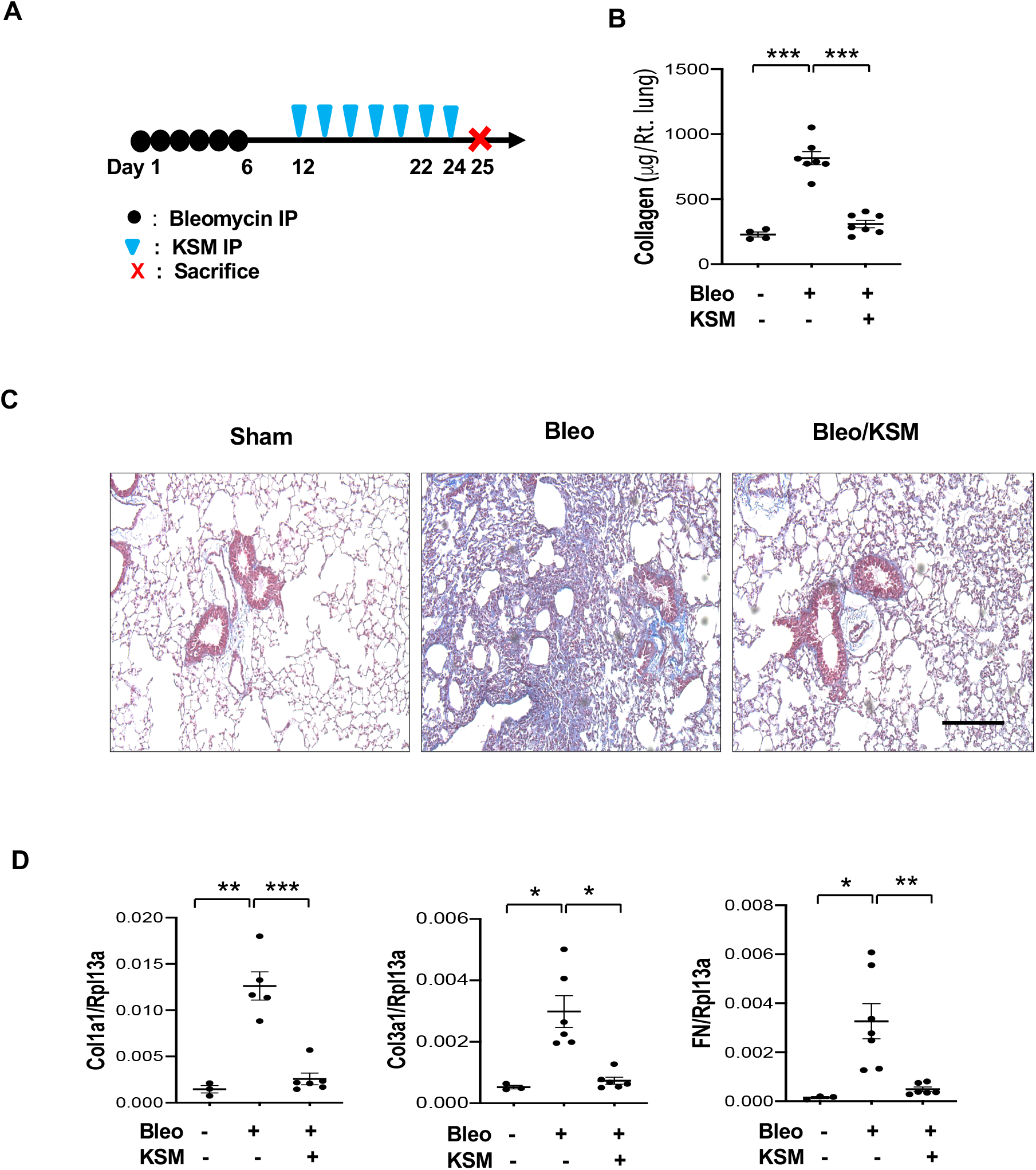
Kasugamycin reduces bleomycin-stimulated pulmonary fibrosis after establishment of fibrosis. 6**-**8 weeks old male mice were challenged with bleomycin (Bleo; i.p, 0.25U/mouse) first 6 days then the mice were treated with Kasugamycin (KSM) (50 mg/kg/mouse, i.p., every other day). (A) Schematic illustration of the protocol employed in this study. (B) Sircol collagen quantitation in the lungs. (C) Representative histology with Mallory’s trichrome staining of the lungs. (D) Real-time qPCR evaluation of collagen I (Col1a1), III (Col3a1) and fibronectin (FN) expression in the lung. Rpl13a was used as an internal control. Each value in panels B and D is from a different animal and the mean±SEM are illustrated. Each value in panels B and D is from a different animal and the mean±SEM are illustrated. \*\**P*<0.05, \*\**P*<0.01, \*\*\**P*<0.001. (One-Way ANOVA with Turkey post hoc test for multiple comparisons). Scale bars=70μm.

### KSM inhibits TGF-β-stimulated fibrosis in the lung

Studies undertaken to see whether KSM has an antifibrotic effect in TGF-β-stimulated pulmonary fibrosis. After induction of TGF-β transgene expression by doxycycline (0.5g/L via drinking water), WT and TGF-β Tg mice were given vehicle (Phosphate Buffered Saline; PBS) or KSM for 2 weeks (every other day, 50mg/kg of KSM or vehicle, i.p) as illustrated in Figure 5A. Collagen accumulation was increased in the lungs of TGF-β Tg mice and KSM treatment significantly reduced TGF-β-induced collagen accumulation in the lung compared to vehicle-treated controls (Fig 5, B and C). We also noted similar changes in the expression of TGF-β stimulated extracellular matrix genes such as Col1a1, Col3a1 and fibronectin (data not shown).

**Figure 5.**
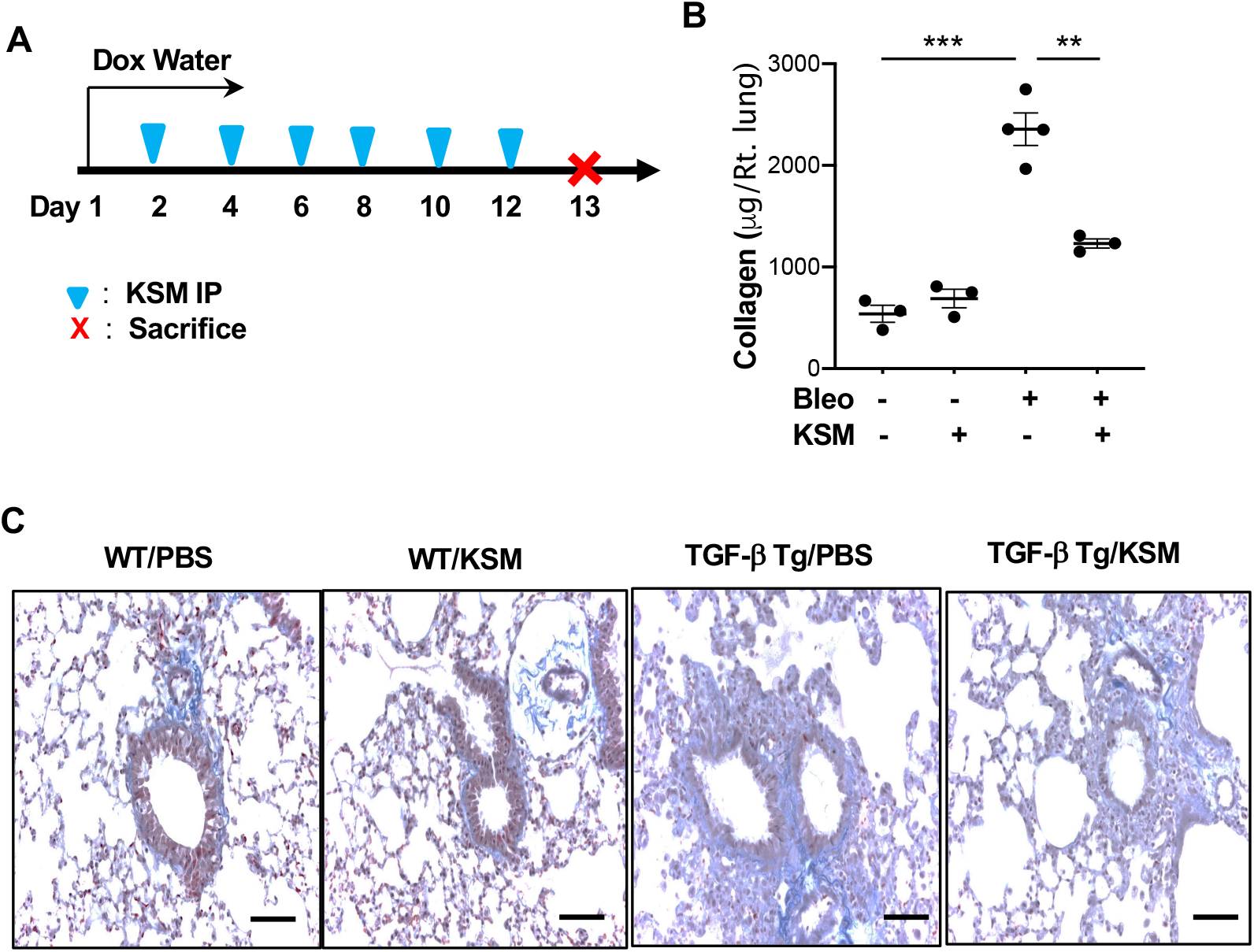
Kasugamycin inhibits TGF-β-stimulated pulmonary fibrosis. 8 weeks old WT and TGF-β Tg male mice were subjected to the evaluations. After transgenic induction of TGF-β by doxycycline, the mice were given vehicle or Kasugamycin (KSM) every other day for 2 weeks (via i.p.) and fibrotic tissue responses in the lung was determined. (A) Schematic illustration of the protocol employed in this study. (B) Sircol collagen quantitation in the lungs. (C) Representative histology with Mallory-trichrome staining of the lungs. Each value in panels B is from a different animal and the mean±SEM are illustrated. \*\**P*<0.01, \*\*\**P*<0.001 (One-Way ANOVA with multiple comparisons). Scale bars =70μm.

### Effect of KSM in profibrotic macrophage activation

Since macrophages are the major cells expressing CHIT1 in IPF lungs, we evaluated the role of CHIT1 and the effect of KSM in specific macrophage activation. In this evaluation, we used primary alveolar macrophages isolated from WT mice and Chit1 null mutant mice (*Chit1*^-/-^). The macrophages were incubated with recombinant (r) IL-4, rTGF-β or rIFN-*γ* for 24 hr in the presence and absence of KSM or vehicle (PBS), then the expression of specific macrophage activation markers was evaluated as we recently described (18). Incubation of alveolar macrophages with rIL-4 (20ng/ml) or rTGF-β (20ng/ml) induced the gene expression of profibrotic activation markers of macrophages (CD206, CD163, CD204, Col1a1, Col3a1) (Figure 6.). The induction of these profibrotic markers was abrogated in the macrophages treated with KSM (250ng/ml) or null mutation of CHIT1 compared to vehicle controls (Figure 6). On the other hand, rIFN-*γ* (10ng/ml)-stimulated inducible nitric oxide synthase (iNOS) expression was not significantly altered with KSM treatment or null mutation of CHIT1 (Figure 6). These studies suggest that CHIT1 plays an essential role in profibrotic macrophage activation and KSM is a potent inhibitor of CHIT1-mediated profibrotic macrophage activation.

**Figure 6.**
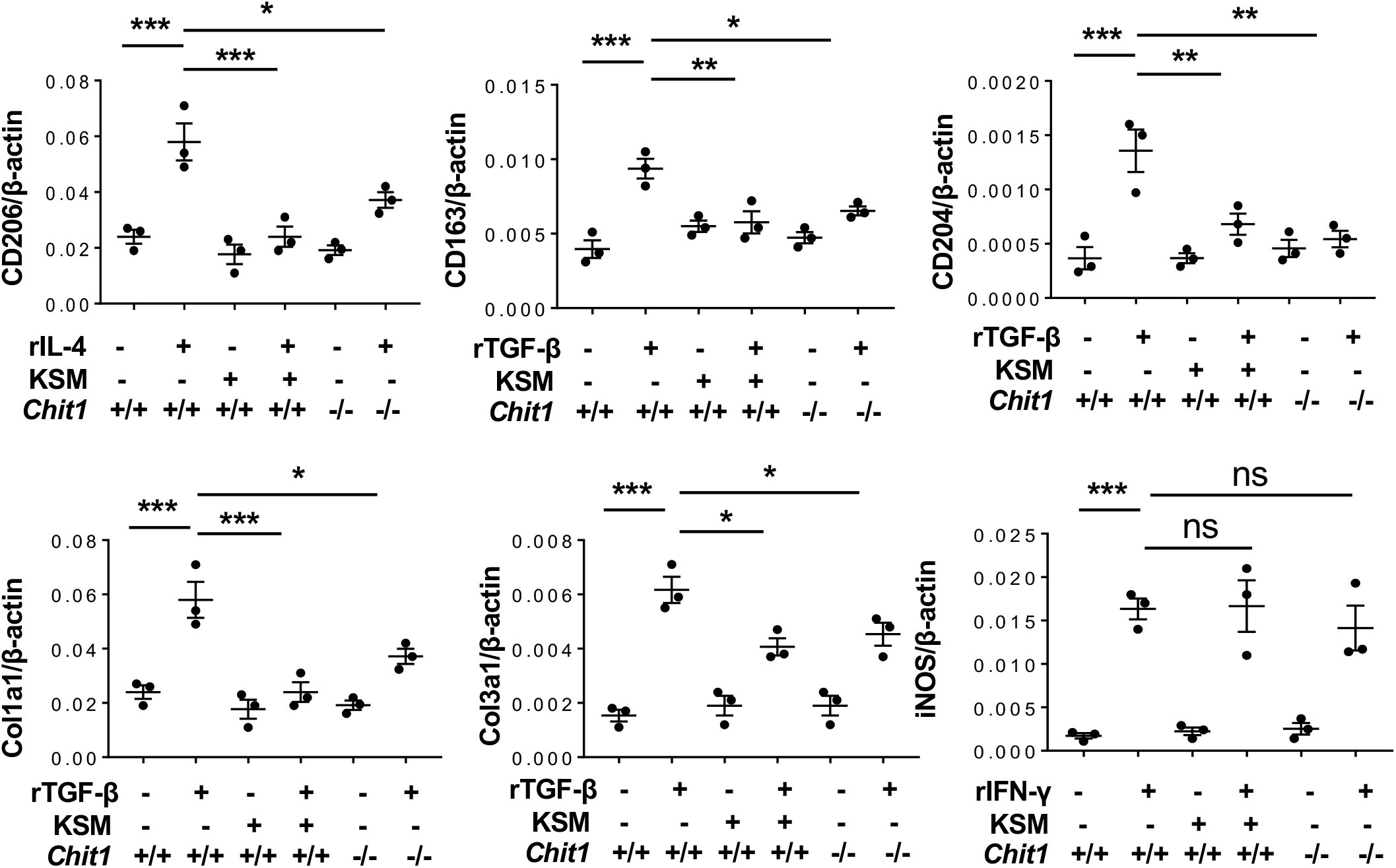
Kasugamycin inhibits profibrotic macrophage activation. Primary alveolar macrophages from WT and *Chit1*^-/-^ mice were stimulated with recombinant (r) IL-4, rTGF-β and rIFN-*γ*, and incubated with (+) and without (-) Kasugamycin (KSM). The mRNA expression of markers associated with macrophage activation was assessed using real time RT-PCR. β-actin was used as an internal control. The values represent the mean±SEM of triplicated evaluations in 2 separate experiments. \**P*<0.05, \*\**P*<0.01, \*\*\**P*<0.001, ns, not significant (One-Way ANOVA with multiple comparisons).

### KSM inhibits CHIT1/TGF-β-stimulated fibroblast proliferation and myofibroblast transformation *in vitro*

Since CHIT1 enhances the TGF-β-stimulated fibroblast proliferation and myofibroblast transformation (5), studies were undertaken to see whether KSM inhibits TGF-β-stimulated fibroblast responses using normal human lung fibroblasts (NHLF) and fibroblasts from a patient with pulmonary fibrosis. In cell proliferation assay, recombinant (r) TGF-β alone or together with rCHIT1 stimulated fibroblast proliferation after 48hr of incubation (Figure 7A, Figure E1). Fibroblast proliferation was significantly reduced with KSM treatment compared to vehicle treated cells (Figure 7A, Figure E1). Similarly, in the cells stimulated with rTGF-β alone or together with rCHIT1 showed enhanced *α*-smooth muscle actin stains and mRNA expression compared to vehicle controls and these increases were abrogated by KSM treatment similarly to Nintedanib (Figure 7, B and C). These studies demonstrated that KSM inhibits TGF-β-stimulated fibroblast proliferation and myofibroblast transformation. Our studies demonstrated significant KSM effects on commercially available human lung fibroblasts derived from both healthy individual and a patient with pulmonary fibrosis. However, considering significant heterogeneity of fibroblasts in fibrotic lungs (19, 20), to generalize these findings, KSM effects need to be further validated using fibroblasts directly derived from multiple patients with pulmonary fibrosis.

**Figure 7.**
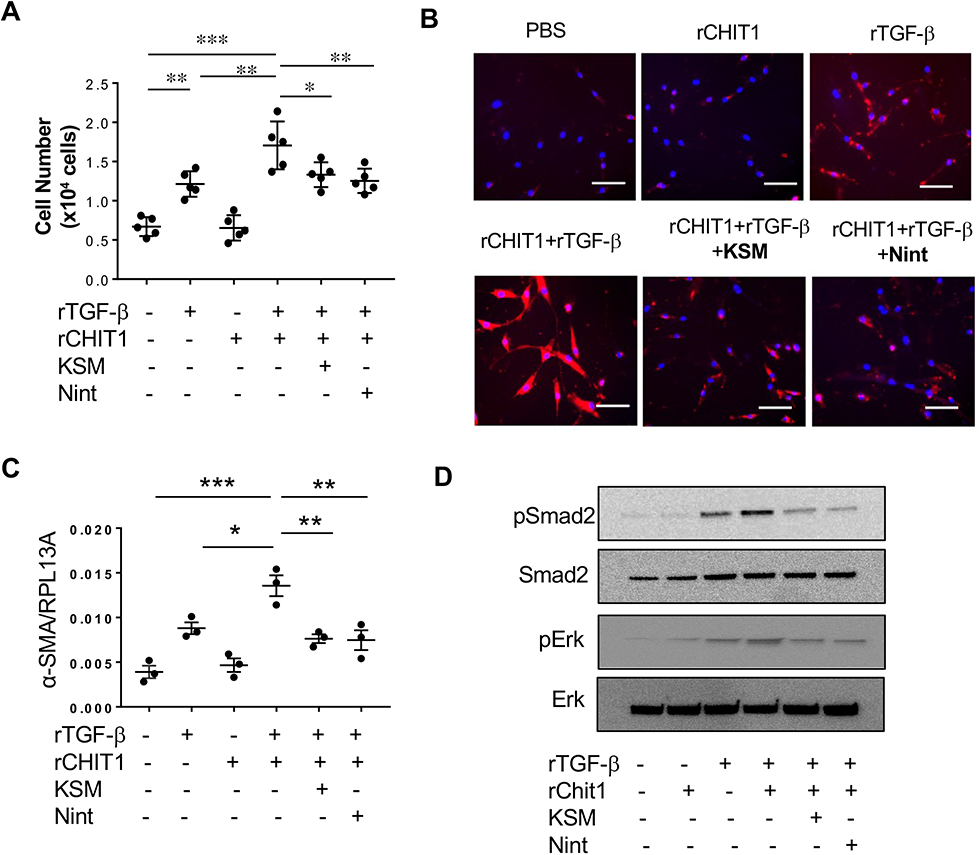
Kasugamycin inhibits CHIT1/TGF-β-stimulated profibrotic fibroblast proliferation, Smad/ERK signaling and myofibroblast transformation. Lung fibroblasts established from pulmonary fibrosis patient (ATCC, PCS-201-020) were stimulated with rTGF-β (10ng/ml) alone or together with rChit1(25ng/ml) for 48 hr. The cells were also incubated with Kasugamycin (KSM) (0.5μM) or Nintedanib (1μM) and fibroblast responses were evaluated. (A) The viable cell numbers were determined after the suspended cells stained with 0.4% trypan blue. (B) *α*-smooth muscle actin IHC to assess myofibroblast transformation. (C) Real-time RT-PCR on the expression of *α*-smooth muscle actin. RPL13A was used as an internal control. (D) Western blots on TGF-β signaling. The values in panels A and C are mean±SEM. Panel D is a representative immunoblot of four separate experiments. \**P*<0.05, \*\**P*<0.01, \*\*\**P*<0.001 (One-Way ANOVA with multiple comparisons). Scale bars=50μm.

### KSM abrogates CHIT1 stimulated TGF-β signaling

Studies were undertaken to see whether KSM modulates the CHIT1-simulated canonical and noncanonical signaling of TGF-β in lung fibroblasts. After NHLF cells were stimulated with rTGF-β or vehicle for 24 hr with KSM and vehicle, then SMADs, MAPK/ERK activation were evaluated by Western blot evaluation. Nintedanib was used as a positive control for inhibition of TGF-β stimulated signaling. As shown in Figure 7D, KSM significantly reduced the CHIT1 and TGF-β activation of SMAD2 and ERK to comparable levels to the cells treated with Nintedanib.

### Role of TGFBRAP1 in bleomycin-stimulated fibrosis and KSM effect on the interaction between CHIT1 and TGFBRAP1

Recently we have identified that TGFBRAP1 is a CHIT1 interacting protein that mediates the CHIT1 effects on TGF-β signaling (5). To determine the specific *in vivo* role of TGFBRAP1 and for further mechanistic characterization of antifibrotic effect of KSM, we generated TGFBRAP1 null mutant mice using Crispr/Cas9 approach (Figure E2). The TGFBRAP1 null mutant mice did not show any apparent abnormal phenotypes in general and demonstrated normal lung development with no sign of inflammation (Figure E2). When WT (*Tgfbrap1*^+/+^) and TGFBRAP1 null mutant mice (*Tgfbrap1*^-/-^) were challenged with bleomycin, collagen accumulation and the expression of collagen genes in the lungs of *Tgfbrap1*^-/-^ mice were significantly decreased compared to WT animals (Figure 8, A and B). The bleomycin-stimulated canonical (Smad2) and non-canonical TGF-β signaling (Erk and Akt) was also significantly reduced in the lungs of *Tgfbrap1*^-/-^ mice compared to WT animals (Figure 8C). To see whether fibroblast response is altered by genetic ablation of TGFBRAP1, we established the primary fibroblasts from the lungs of WT and *Tgfbrap1*^-/-^ mice. As shown in Figure 8D and 8E, the rTGF-β alone or rTGF-β and rChit1 co-stimulation significantly enhanced lung fibroblast proliferation and myofibroblast transformation in WT fibroblasts, but these responses were abrogated in the lung fibroblasts of *Tgfbrap1*^-/-^ mice. Lastly, studies were performed to evaluate the effect of KSM on the interaction of CHIT1 and TGFBRAP1 using co-immunoprecipitation (Co-IP) and immunoblot (IB) assays in HEK293 cells co-transfected with human CHIT1 and TGFBRAP1 expressing plasmids. As shown in Figure 8F, coimmunoprecipitation of CHIT1 and TGFBRAP1 support a physical association between CHIT1 and TGFBRAP1 as we previously reported (5). The treatment of KSM on these co-transfected cells significantly reduced the co-precipitation of TGFBRAP1 and CHIT1 (Figure 8F), suggesting that KSM interferes the physical association between CHIT1 and TGFBRAP1. When viewed in combination, these studies demonstrated that TGFBRAP1 plays an essential role in bleomycin- or CHIT1/TGF-β-stimulated fibrotic tissue and cellular responses and that KSM inhibits CHIT1 association with TGFBRAP1 and SMAD4 complex.

**Figure 8.**
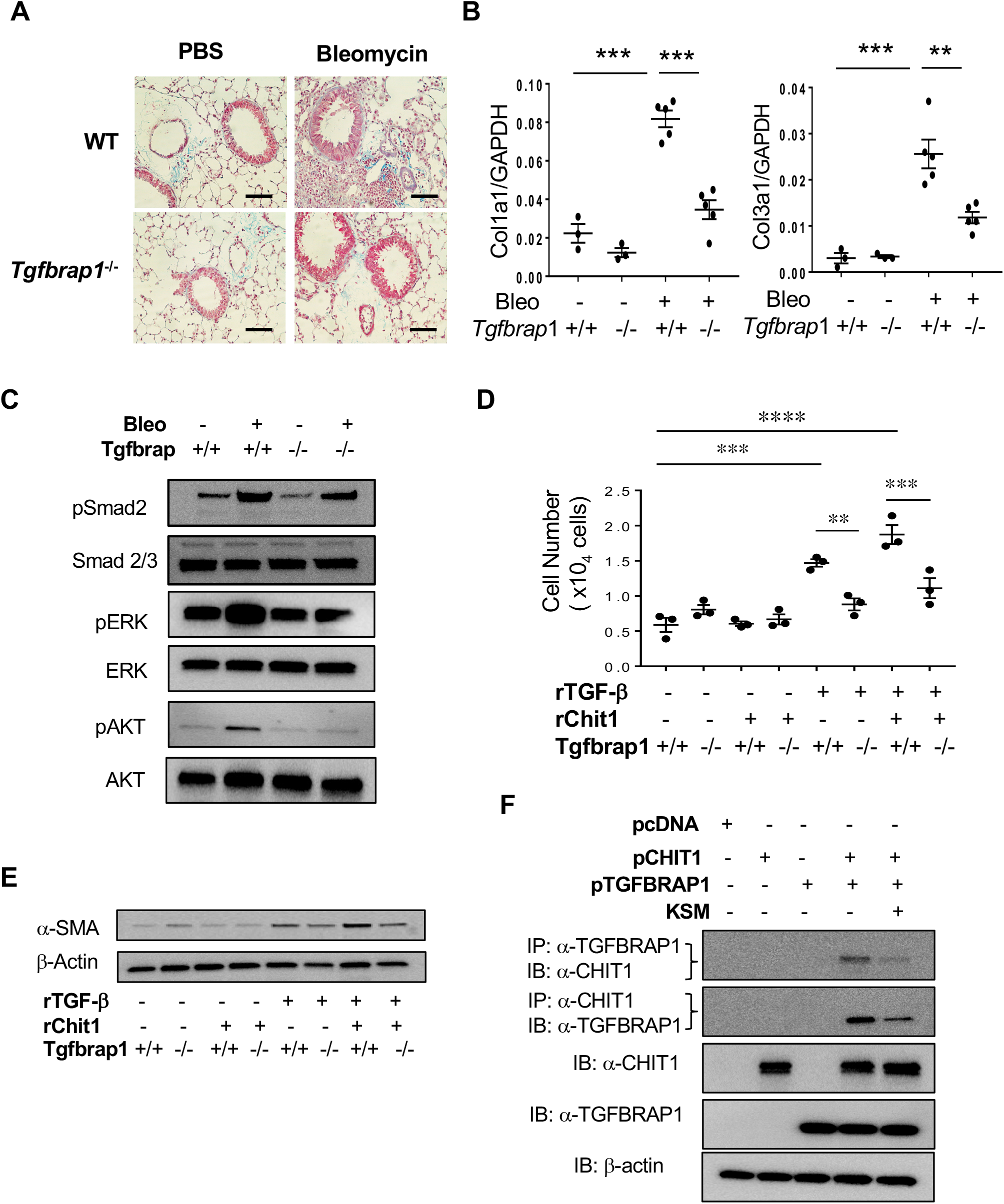
Role of TGFBRAP1 in bleomycin-stimulated lung fibrosis and CHIT1 augmentation of TGF-β1-stimulated fibroblast response and the effects of Kasugamycin on CHIT1 and TGFBRAP1 interaction. 8 weeks old WT and *Tgfbrap1*^-/-^ mice (-/-) were challenged with bleomycin (Bleo; i.p, 0.25U/mouse) then the mice were sacrificed on day 25 and subjected to the evaluation. (A) Representative histology with Mallory’s-trichrome staining of the lungs. (B) Real-time PCR evaluation of collagen I (Col1a1) and III (Col3a1) expression in the lung. (C) Western blot evaluations of TGF-β signaling in the lung. (D-E) Fibroblasts established from WT (+/+) and *Tgfbrap1*^-/-^ (-/-) were stimulated with rTGF-β (10ng/ml) and rCHIT1 (5μM) and fibroblast proliferation and myofibroblast transformation (*α*-smooth muscle actin (SMA) expression) were evaluated by cell proliferation assay (D) and Western blot evaluations (E), respectively. (F) HEK293 cells were transfected with CHIT1 and TGFBRAP1 cDNAs and the cell lysates were subjected to the co-immunoprecipitation (IP) and immunoblot (IB) assays using and *α*-CHIT1, and *α*-TGFBRAP1 antibodies in the presence and absence of Kasugamycin (KSM). Each value in panels B and D is from a different animal and the mean±SEM are illustrated. The panel D is a representative of 2 separate experiments. Panels C, E, and F are representative immunoblots of four separate experiments. \*\**P*<0.01, \*\*\**P*<0.001, \*\*\*\**P*<0.0001 (One-Way ANOVA with multiple comparisons). Scale bars=70μm

## DISCUSSION

Pulmonary fibrosis is a devastating lung disease that affects more than 200,000 people in the US alone. Idiopathic pulmonary fibrosis (IPF) is the most common form of pulmonary fibrosis (PF) and has a poor life expectancy (two to three years of median survival) after diagnosis (21, 22). Currently, there are two FDA approved therapeutic drugs for PF: Pirfenidone (Esbriet^®^) and Nintedanib (Ofev or Vargatef^®^). These drugs provide substantial benefits by slowing disease progression, but they do not offer symptomatic relief or improvement in quality of life and are associated with various side effects and prohibitively high cost (23–25). Additionally, available treatments do not reverse lung damage incurred during fibrotic progression, necessitating early treatments prior to the destruction of normal lung architecture. The only curative treatment for PF is a high-risk lung transplant and lifelong anti-rejection treatments. However, lung transplants are associated with extreme morbidity and recipients often succumb to fibrotic rejection within 5-10 years of the transplant. Therefore, new drug or methods of prevention and treatment of PF are critically needed.

In the previous studies, we identified CHIT1 is significantly implicated in the pathogenesis of pulmonary fibrosis. Specifically, we demonstrated a significant increase in the expression of CHIT1 in the lungs of patients with scleroderma associated interstitial lung disease (SSc-ILD) (4). We also demonstrated increased levels of CHIT1 activity in the patient with SSc-ILD that were inversely correlated with lung function and overall progression free survival (4, 5). In current study, we further identified that expression of CHIT1 is significantly increased in the lungs of IPF patients compared to normal lungs and alveolar macrophages, airway epithelial cells, and fibroblast-like parenchymal cells are the major cells expressing CHIT1. Genetic ablation of CHIT1 significantly reduced collagen accumulation in the lungs of multiple animal models of fibrosis, while lung-specific overexpression of CHIT1 enhanced bleomycin- or TGF-β-stimulated fibrosis (4, 5). These studies strongly support an essential role of CHIT1 in the pathogenesis of pulmonary fibrosis. Thus, we believe that CHIT1 inhibition would be an effective therapeutic strategy for pulmonary fibrosis.

In an effort to identify CHIT1 inhibitor(s) to be used for antifibrotic agent(s), we performed high throughput small molecule screening using established libraries. Intriguingly, these studies identified KSM, an aminoglycoside antibiotic, is a strong and specific CHIT1 inhibitor. In validation studies using animal models of pulmonary fibrosis, we were able to appreciate impressive preventive as well as therapeutic effects of KSM that significantly reduced bleomycin- or TGF-β-stimulated fibrotic responses in the lung. In this study, we further validated antifibrotic activity of KSM as a CHIT1 inhibitor and determined the potential anti-fibrotic mechanism of KSM in the pathogenesis of pulmonary fibrosis.

KSM is a natural compound produced by *Streptomyces kasugaensis* originally isolated from soil at Kasuga Shrine in Nara, Japan (26). Since it is an aminoglycoside antibiotic, multiple groups investigated KSM for possible clinical utility following its introduction. Initial studies suggested that KSM was effective against *Pseudomonas* (12, 27). However, subsequent *in vitro* susceptibility testing yielded mixed results, and small trials using Kasugamycin as an antibiotic against *Pseudomonas* infections in animals and humans were disappointing (28–30). With these reasons, KSM has not been used as an antibiotic for the treatment of human diseases. However, it is still widely used in the agricultural realm to control fungal and bacterial infections in fruits, rice and other crops especially in organic farming and its minimal or no toxicity to humans and animals are well established (10, 11).

The mechanism of anti-chitinase activity of KSM has not been determined yet. Unlike other aminoglycosides, KSM has a unique structure: it is composed of two sugars, a D-chiro-inositol moiety and a kasugamine moiety (2,4-diamino-2,3,4,6-tetradeoxy-Darabino-hexose), with a carboxy-imino-methyl group (31). This structural character of KSM as a carbohydrate compound may confer its ability to bind with CHIT1 as a substrate like oligo-chitin (32). Anti-chitinase activity of oligo-chitin has been demonstrated (32). If this is the case, KSM can occupy the catalytic or chitin binding domains of the CHIT1 and block the chitin-degrading enzyme activity of CHIT1. However, the exact nature of physical binding between CHIT1 and KSM remains to be determined. In addition, KSM binding to CHIT1 could also affect the interactions between CHIT1 and other molecules responsible for profibrotic activities of CHIT1. This notion led us to investigate the role of TGFBRAP1, a recently defined CHIT1 interacting molecule (5), and the KSM effect on the interaction between CHIT1 and TGFBRAP1 as discussed below.

IPF, a prototypic fibrotic disorder, is a progressive chronic lung disease characterized by epithelial damage, fibroproliferative matrix deposition and parenchymal remodeling (33–35). TGF-β is believed to play an essential role in this dysregulation because it is expressed in an exaggerated fashion in IPF where, in contrast to controls, a sizable percentage is biologically active (36–38). However, the mechanisms through which TGF-β controls fibrotic tissue responses in the lungs are not sufficiently determined. In the previous studies (5), we have demonstrated that CHIT1 enhances pulmonary fibrosis by augmenting TGF-β signaling and effector function through interaction with TGFBRAP1, a chaperone for SMAD4, that recruit SMAD4 to the vicinity of the receptor complex to facilitate its interaction with receptor-regulated SMADs, such as SMAD2/3 (6). In this study, we have demonstrated a significant role of TGFBRAP1 in the pathogenesis of pulmonary fibrosis by generating and characterization of TGFBRAP1 null mutant mice (*Tgfbrap1*^-/-^). It is also of interesting to note that null mutation of TGFBRAP1 phenocopied the effects of KSM on pulmonary fibrosis *in vivo* and fibroblasts responses *in vitro*. The immunoprecipitation and immunoblot assays further confirmed that CHIT1 physically interacts with TGFBRAP1 and KSM inhibits the association between these two molecules. Since TGFBRAP1 is a known as a chaperon of SMAD4 (6), we can readily envision that disruption of CHIT1 and TGFBRAP1 by KSM treatment could result into less Smad4 recruitment as a mechanism of KSM uses to inhibit TGF-β signaling. However, the impact of KSM on the interaction of TGFBRAP1 and SMAD4 in the presence and absence of CHIT1 remains to be determined in future studies. When viewed in combination, these studies suggest a possibility that KSM inhibits CHIT1 mediated TGF-β signaling and effector function through regulation of interaction between CHIT1 and TGFBRAP1, at least in part.

In summary, we demonstrated that CHIT1 expression is dysregulated in the lungs of patients with pulmonary fibrosis. Based on the prior studies supporting CHIT1 as a therapeutic target of pulmonary fibrosis, we sought druggable CHIT1 inhibitor through high throughput small molecule screening. In this study, we have identified that KSM is a novel CHIT1 inhibitor that has a strong antifibrotic activity against bleomycin- or TGF-β-stimulated pulmonary fibrosis potentially through inhibition of CHIT1 and TGFBRAP1 interaction. These studies suggest that KSM can be further developed as a new CHIT1-based antifibrotic agent that effectively and safely blocks the development and or progression of pulmonary fibrosis.

## MATERIALS AND METHODS

### Mice

C57BL/6 mice were purchased from the Jackson Laboratory (Bar Harbor, ME, USA) and were housed at Brown University animal facilities. *Chit1*^-/-^ mice were generated and characterized as previously described (5). Tgfbrap1 null mutant mice (*Tgfbrap1*^-/-^) on C57 BL/6 background were generated at transgenic facility at Brown University using *Crisp/Cas9* approach and maintained in our laboratory as previously described (39). TGF-β Tg mice are lung-specific inducible transgenic mice and transgene expression was induced by drinking water containing Doxycycline (0.5g/L)(39). The *Tgfbrap1*^-/-^ mice did not show any apparent abnormal phenotypes or developmental issues. All murine procedures were approved by Brown Institutional Animal Care and Use Committees.

### High throughput small molecule library screening

Using chitinase activity as a quantitative read-out to efficiently identify CHIT1 inhibitors, a total of 7670 small molecules were screened through the Yale Center for Molecular Discovery using a collection of libraries. Please see detailed screening methods in the online supplement. They include Microsource GenePlus, Microsource Pharmakon, Enzo640 FDA-approved drugs and Yale Procured Drugs Collections, Selleck Chem Kinase Inhibitors, NIH Clinical Collections, NCI Diversity Set 2, NCI Oncology Set 2, Microsource Natural Products, and ENZO Protease Inhibitors. In this screening process, inhibition of CHIT1 enzyme activity was measured using fluorogenic substrate 4-methylumbelliferyl β-D-N,N′,N′′-triacetylchitotrioside (4MU-GlcNAc3; Sigma-Aldrich, St Louis, MO) following hydrolysis measuring liberated fluorescent 4-in McIlvain buffer (100 mmol/l citric acid and 200 mmol/l sodium phosphate [pH 5.2]) as described previously (40). Briefly, in a final volume of 50 μl, 2 nM of recombinant CHIT1 (R&D Systems. Cat # 3559-GH-010) was incubated with 20 μM substrate in McIlvain buffer (100 mM citric acid, 200 mM sodium phosphate [pH 5.2]) containing 0.1 mg/ml BSA, for 30 min at 37°C in the presence of specific CHIT1 inhibitor candidate. After the addition of 25 μl of 3 M glycine-NaOH (pH 10.3), the fluorescence of the liberated 4MU was quantified using high throughput plate reader with excitation and emission wavelengths of 360 nm and 460 nm, respectively. The activity of CHIT1 inhibition of candidate drug was expressed as % effect (% inhibition) relative to pentoxifylline (200μM), a known CHIT1 inhibitor (7).

### *In vivo* evaluation of KSM effects in animal models of lung fibrosis

*in vivo* effects of KSM in pulmonary fibrosis were assessed using bleomycin model of lung fibrosis and lung-specific TGF-β transgenic mice established in our laboratory were used (4, 39). The effective in vivo dose and route of delivery (12.5-100mg/kg/mouse, i.p injection, every other day) were determined based on previously reported animal studies (41) and preliminary studies that demonstrated sustained anti-CHIT1 activity in the lung until 2 days after KSM administration (Supplemental Figure E3).

### Statistical analysis

Statistical evaluations were undertaken with GraphPad Prism software. As appropriate, groups were compared with 2-tailed Student’s *t* test or with nonparametric Mann-Whitney *U* test. Values are expressed as mean ± SEM. One-way ANOVA or nonparametric Kruskal-Wallis test were used for multiple group comparisons. Statistical significance was defined as a level of *P* < 0.05.

Please *see* other methodologies employed in this study in the online data supplement. They include cell culture, histologic analysis, Real-time qRT-PCR, Sircol collagen assay, Western blotting, double label fluorescent Immunohistochemistry, fibroblast proliferation assay, establishment of primary fibroblasts, Isolation of alveolar macrophages from the lungs of mice and assessments macrophage activation, and Co-immunoprecipitation and immunoblot assays. All the Western blot evaluations were quantitated using ImageJ program and provided as a supplemental information (Figure E4).

## Impact of this research on clinical medicine and basic science

Idiopathic pulmonary fibrosis is a devastating lung disease with few therapeutic options. This study identified Kasugamycin as a chitinase 1 inhibitor with strong antifibrotic activity that can be developed as an effective and safe therapeutic drug for the patients with pulmonary fibrosis. This study also provides a novel mechanism that Kasugamycin uses to control fibrotic tissue and cellular responses in the lung.

## Author Contributions

Conception and design: JHL, CML, CGL; Generation of experimental resources and data collection: JHL, CML, MOK, JWP, SK, BA, ELH, XP, JoHL; Analysis and interpretation: JHL, CML, JAE, CGL; Drafting the manuscript for important intellectual content: JHL, JAE, CGL.

## Acknowledgements (Grants and Other Supports)

This work was supported by National Institute of Health (NIH) grants PO1 HL114501(JAE) and R01 HL115813 and RO1 HL155558 (CGL) from NHLBI and P20 GM103652 from NIGMS (CML). This work was also partly supported by sponsored research grant (GR5290824) (CGL) from Corestem Inc. and Brown Biomedical Innovations to Impact (BBII) grant (CGL) from Brown University and W81XWH2210041 (CGL) from Peer Reviewed Medical Research Program of Department of Defense.

## Competing Interests

JAE is a cofounder of Elkurt Therapeutics and is a founder of, stock holder of, and serves on the Scientific Advisory Board for Ocean Biomedical, Inc., which develops inhibitors of 18 glycosyl hydrolases as therapeutics. The other authors have declared that no conflict of interest exists.

This article has an online data supplement, which is accessible from this issue’s table of content online at www.atsjournals.org.

