## Supplementary material for "Kasugamycin is a novel chitinase 1 inhibitor with strong antifibrotic effects on pulmonary fibrosis": Online Data Supplement

**Supplementary Methods:**

**Cell culture**

Normal human lung fibroblast cells (NHLF; CC2512) and primary lung fibroblasts established from patient with lung fibrosis (PCS-201-020) were purchased from Lonza (Allendale, NJ) and American Type Culture Collection (ATCC, Manassas, VA), respectively. Human embryonic kidney 293 cells (HEK293; CRL1573) were obtained from ATCC. NHLF cells were cultured in fibroblast growth medium (FGM; Lonza, CC-2512B) at 37°C in 5% $CO_2$. Primary lung fibroblasts from fibrosis patients were cultured using fibroblast basal medium supplemented with fibroblast growth kit as recommended by ATCC. HEK293 cells were maintained in Dulbecco's modified Eagle's medium (DMEM) supplemented with 10% fetal bovine serum (FBS) and 100 U/ml penicillin and 100 µg/ml streptomycin at 37°C in 5% $CO_2$ incubator. Both primary cells of lung fibroblasts were used within 10 passages according to the manufacturer's recommendation.

**Histologic Analysis**

The lungs were removed en bloc, inflated at 25 cm pressure with neutral buffered 10% formalin, fixed in 10% formalin, embedded in paraffin, sectioned, and stained. Hematoxylin and eosin and Mallory's trichrome stains were performed in the pathology core at Brown University.

**Sircol collagen assay**

Animals were anesthetized, a median sternotomy was performed, and right heart perfusion was completed with calcium and magnesium-free PBS. The heart and lungs were then removed. The right lung was frozen in liquid nitrogen and stored at −80 °C until they were used. Collagen content was determined by quantifying total soluble collagen using the Sircol collagen assay kit (Biocolor, Accurate Chemical & Scientific Co., Westbury, NY) according to the instructions of the manufacturer.

**Real-time RT-PCR**

mRNA levels were assessed using real-time RT-PCR assays. In these assays, total cellular RNA from lungs or other mouse tissues were obtained using trizol reagent (GIBCO BRL) and RNAeasy mini Kit (Qiagen, Frederick, MD) according to the manufacturer's instructions. The information on the primers used in this study is included in supplementary Table E2.

**Western blotting (Immunoblotting)**

25 µg lung or cell lysates were subjected to immunoblot analysis using antibodies against $\alpha$-smooth muscle actin ($\alpha$-SMA), Phosphorylated(p) ERK (pERK), total ERK(ERK), Phosphorylated(p) AKT (pAKT), total AKT(AKT). These samples were gel fractionated, transferred to membranes, and evaluated as described previously by our laboratory (1).

**Isolation of alveolar macrophages from the lungs of mice**

Mice were sacrificed using intraperitoneal (i.p.) ketamine/xylazine injection, and the trachea was cannulated and perfused with two 0.9 ml aliquots of cold saline. The cellular contents were resuspended into RPMI1640 media with 10% FBS and 1X Penicillin/Streptomycin and were plated 10 $cm^2$ tissue flasks at 37°C/5% CO2. After 1 day, nonadherent cells were discarded and medial was changed. Purity of macrophages (>95 %) was confirmed via flow cytometry.

**Immunohistochemical (IHC) evaluation**

To evaluate the expression of CHIT1 in the lung and $\alpha$-SMA in lung fibroblasts, IHC was undertaken with a modification of procedures described previously by our laboratory (2). First, slides were deparaffinized in xylene, rehydrated in an ethanol gradient, and washed in PBS. To unmask antigens, slides were placed for 20 min at high temperature under high pressure in citrate buffer (10 mM sodium citrate, 0.05% Tween 20, pH 6). Tissue sections were then blocked with a non-serum protein-blocking reagent (DakoCytomation Inc., Mississauga, ON, Canada) for 1 hr at room temperature and incubated with primary antibodies (anti-CHIT1 (1/50 dilution, AF-3559, R&D Systems); anti-$\alpha$-smooth muscle actin (1/50 dilution, A-2547 Sigma-Aldrich St Louis, MO) for 60 min at room temperature in a humid chamber. Substitution of the primary antibody with PBS served as a negative control. The slides were then washed in PBST (0.01 % Tween 20), incubated with secondary antibodies conjugated with horseradish peroxidase (Cell signaling MA, USA). Reaction products were developed using diaminobenzidine (DAB) containing 0.3% hydrogen peroxide up to 5 min.

**Double label fluorescent Immunohistochemistry**

Formalin-fixed paraffin embedded (FFPE) lung tissue blocks were serially sectioned at 5 µm thickness and mounted on glass slides. After deparaffinization and dehydration, heat-induced epitope retrieval was performed by boiling the samples in a steamer for 30 minutes in antigen unmasking solution (Abcam, antigen retrieval buffer, 100x citrate buffer pH:6.0). To prevent nonspecific protein binding, all sections were blocked in a ready-to-use serum free protein blocking solution (Dako/Agilent, Santa Clara, CA) for 10 minutes at room temperature. The sections were then incubated with primary antibodies ($\alpha$-CHIT1 (Sigma-Aldrich, HPA010575), $\alpha$-F4/80 (Santa Cruz Biotechnology, sc-377009), $\alpha$-CC10 (Santa Cruz Biotechnology, sc-365992), $\alpha$-SPC (Santa Cruz, sc-518029)) overnight at 4°C. After three washings, fluorescence-labeled secondary antibodies were incubated for 1 hour at room temperature. The sections were then counterstained with DAPI and cover slips were added.

**Viable cell count and WST-1 assay**

The viable cell numbers were determined after the suspended cells stained with 0.4% trypan blue according to previously reported procedures (3). Cells grown in a 96-well tissue culture plate are incubated with the WST-1 reagent (5015944001. Sigma-Aldrich St Louis, MO) in a ratio of 1:10 for 2 hours. After this incubation period, the formazan dye formed is quantitated with a scanning multi-well spectrophotometer (ELISA reader) at 450nM.

**Assessments of macrophage activation**

Alveolar macrophages were incubated in RPMI 1640 plus 10% FCS for 48 h with or without 5 ng/ml IL-4, 10 ng/ml rTGF-$\beta$, or 5 ng/ml of IFN-$\gamma$. Gene expression associated with macrophage activation was evaluated by real-time RT-PCR as described previously (4)

**Establishment of primary lung fibroblasts**

According to the reported protocol (5), ex vivo lung was first chopped into small pieces, then incubated in Liberase Blendzyme solution (grade III, 0.14 Wünsch U/ml, Roche) for 30 minutes under a tube rotator in a 37°C incubator. The suspension was filtered through a 70μm nylon mesh and was centrifuged at 400g for 3 minutes. The cell pellet was resuspended into DMEM/F12 media with 15% FBS, 1× antibiotic/antimycotic and plated in a 10cm$^2$ tissue culture dish at 37°C/5% CO2. Fourteen days after the beginning of the cell isolation, it was changed into EMEM with 15% FBS and 1× penicillin/streptomycin.

**Co-Immunoprecipitation (Co-IP) and immunoblot (IB) analysis**

To assess interactions between CHIT1 and Tgfbrap1, HEK293 cells were co-transfected with TGFBRAP1 and CHIT1 plasmids (pcDNA3.1). Lysates from these cells were subjected to immunoprecipitation using anti-hCHIT1 mouse monoclonal antibody (R&D systems, AF3559) and anti-hTGFBRAP1 (Sigma-Aldrich, HPA038397). The Catch and Release V2.0 (Reversible Immunoprecipitation System, EMD Millipore) kit was used for Co-IP according to the manufacturer's instruction. The precipitates were then evaluated by immunoblotting with antibodies against CHIT1 (R&D systems), TGFBRAP1 (Sigma-Aldrich) and SMAD4 (Cell Signaling, 38454).

### Supplementary Tables

**Table E1.** List of small molecules with anti-Chit1 activity identified in screening of small molecules.

| Supplier | Supplier ID | Drug Name | % EFFECT (relative to 200μM pentoxifylline) |
|---|---|---|---|
| NCI | NSC51349 | 5,7-dimethoxy-2-pyridin-3-ylchromen-4-one | 131.80 |
| SelleckChem | S1219 | YM201636 | 125.99 |
| NCI | NSC60340 | N(1), N(4)-Bis(4-(4,5-dihydro-1H-imidazol-2-yl)phenyl)-2-(hydroxy(oxido)amino)terephthalamide | 122.71 |
| Microsource | 01505038 | Kasugamycin Hydrochloride | 114.82 |
| SelleckChem | S1116 | Palbociclib (PD-0332991) HCl | 107.17 |
| SelleckChem | S2183 | BGJ398 (NVP-BGJ398) | 104.94 |
| Microsource | 01500861 | Coralyne Chloride | 104.10 |
| NCI | NSC260594 | Benzamide, 4-[(1-methyl-6-nitro-4(1H)-quinolinylidene)amino]-N-[4-[(1-methyl-4(1H)-pyridinylidene)amino]phenyl]- Benzamide, 4-[(1-methyl-6-nitro-4(1H)-quinolinylidene)amino]-N-[4-[(1-methyl-4(1H)-pyridinylidene]amino]phenyl]- | 100.66 |
| NCC | SAM001246893 | Pirenperone (CPD000058507) | 90.45 |
| NCI | NSC69187 | 9-Methoxyellipticine Ellipticine, | 90.23 |
| SelleckChem | S2806 | CEP-33779 | 87.98 |
| NCC | SAM001246706 | Pemoline (CPD000238142) | 88.46 |
| NCI | NSC279836 | Mitoxantrone | 89.40 |
| NCI | NSC36758 | Toluidine Blue | 87.98 |
| SelleckChem | S2727 | Dacomitinib (PF299804, PF299) | 84.90 |
| SelleckChem | S2658 | GSK2126458 (GSK458) | 84.57 |
| NCI | NSC134159 | 4-[2-(5,6-Dioxopyridin-3-yl) hydrazinyl] benzenesulfonamide | 85.06 |
| SelleckChem | S1014 | Bosutinib (SKI-606) | 79.73 |
| SelleckChem | S2774 | MK-2461 | 76.04 |
| NCC | SAM001246768 | Doxorubicin Hydrochloride (CPD000058570) | 73.38 |
| NCI | NSC88882 | 1-Hydroxyphenazine | 72.00 |
| SelleckChem | S2759 | CUDC-907 | 67.28 |
| NCI | NSC7218 | 3,6-Diacetamidoacridine 3 | 65.64 |
| SelleckChem | S2736 | TG101348 (SAR302503); Fedratinib | 65.49 |
| SelleckChem | S1113 | GSK690693 | 64.43 |
| Microsource | 01500872 | Palmatine Chloride | 66.04 |
| NCI | NSC60339 | Wander | 64.72 |
| Microsource | 01500223 | Daunorubicin | 63.85 |

| | | | |
|---|---|---|---|
| NCC | SAM001247045 | Trazodone Hydrochloride (CPD000058520) | 62.95 |
| NCI | NSC353263 | Pyridinyl Molecules | 63.61 |
| SelleckChem | S2692 | TG101209 | 59.07 |
| SelleckChem | S1011 | Afatinib (BIBW2992) | 59.66 |
| NCI | NSC317605 | 11H-Indolo[3,2-c] quinolin-9-amine, 3-chloro-N,N-diethyl-8-methoxy- | 60.48 |
| NCC | SAM001246559 | Epirubicin Hydrochloride CPD000466308) | 59.13 |
| NCI | NSC157966 | Lysergic acid methyl ester | 58.94 |
| NCC | SAM001246796 | Raltitrexed (CPD000469217) | 60.25 |
| NCC | SAM001246651 | Topotecan Hcl (Cpd000466344) | 56.24 |
| SelleckChem | S1147 | Barasertib (AZD1152-HQPA) | 52.46 |
| SelleckChem | S2150 | Neratinib (HKI-272) | 52.37 |
| SelleckChem | S2924 | CHIR-99021 HCl | 52.25 |
| SelleckChem | S2842 | SAR131675 | 51.75 |
| NCI | NSC308848 | 3-Amino-N-(2-diethylaminoethyl)-1,8-naphthalimide | 50.94 |

NCI, National Cancer Institute; NCC, National Cancer Center

**Table E2**. Sequences of RT-PCR primers used in this study

| Gene | Sequence (5'to 3') |
|---|---|
| Col1a1-S | tgaacgtgaccaaaaaccaa |
| Col1a1-AS | gcagaaaaggcagcattagg |
| Col3a1-S | ggagcccctggactaatagg |
| Col3a1-AS | atccatctttgccatcttcg |
| FBN-S | aatggaaaaggggaatggac |
| FBN-AS | ctcggttgtccttcttgctc |
| β-actin-S | ggctgtattcccctccatcg |
| β-actin-AS | ccagttggtaacaatgccatgt |
| RPL13a-S | aggggcaggttctggtattg |
| RPL13a-AS | tgttgatgccttcacagcgt |
| CD206-S | caggtgtgggctcaggtga |
| CD206-AS | tgtggtgagctgaaaggtga |
| CD163-S | tccacacgtccagaacagtc |
| CD163-AS | ccttggaaacagagacaggc |
| iNOS -S | cagaggacccagagacaagc |
| iNOS -AS | tgctgaaacatttcctgtgc |
| CD204-S | ctggacaaactggtccacct |
| CD204-AS | tccccttctctcccttttgt |
| SMA-S | gaggcaccactgaaccctaa |
| SMA-AS | catctccagagtccagcaca |
| TGFBRAP-1 -S | ataggctgctggtgctctgt |
| TGFBRAP-1 -AS | tccttgacaatctgcactcg |
| GAPDH-S | catcactgccacccagaagactg |
| GAPDH-AS | atgccagtgagcttcccgttcag |

### Online Data Supplemental Figures

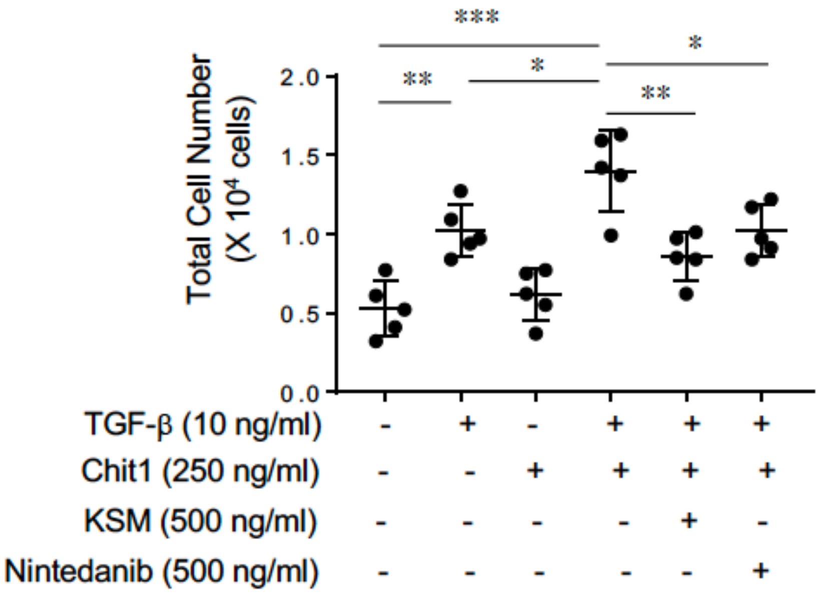


**Figure E1. Kasugamycin inhibits CHIT1/TGF-β-stimulated normal lung fibroblast proliferation**. Normal human lung fibroblasts (NHLF; CC2512, Lonza) were stimulated with rTGF-β (10ng/ml) alone or together with rChit1(25ng/ml) for 24 hr. The cells were also incubated with Kasugamycin (KSM) (0.5μM) or Nintedanib (1μM) and fibroblast responses were evaluated. The viable cell numbers were determined after the suspended cells stained with 0.4% trypan blue. The values are mean±SEM. *$P$<0.05. **$P$<0.01, ***$P$<0.001 (One way ANOVA, multiple comparisons).

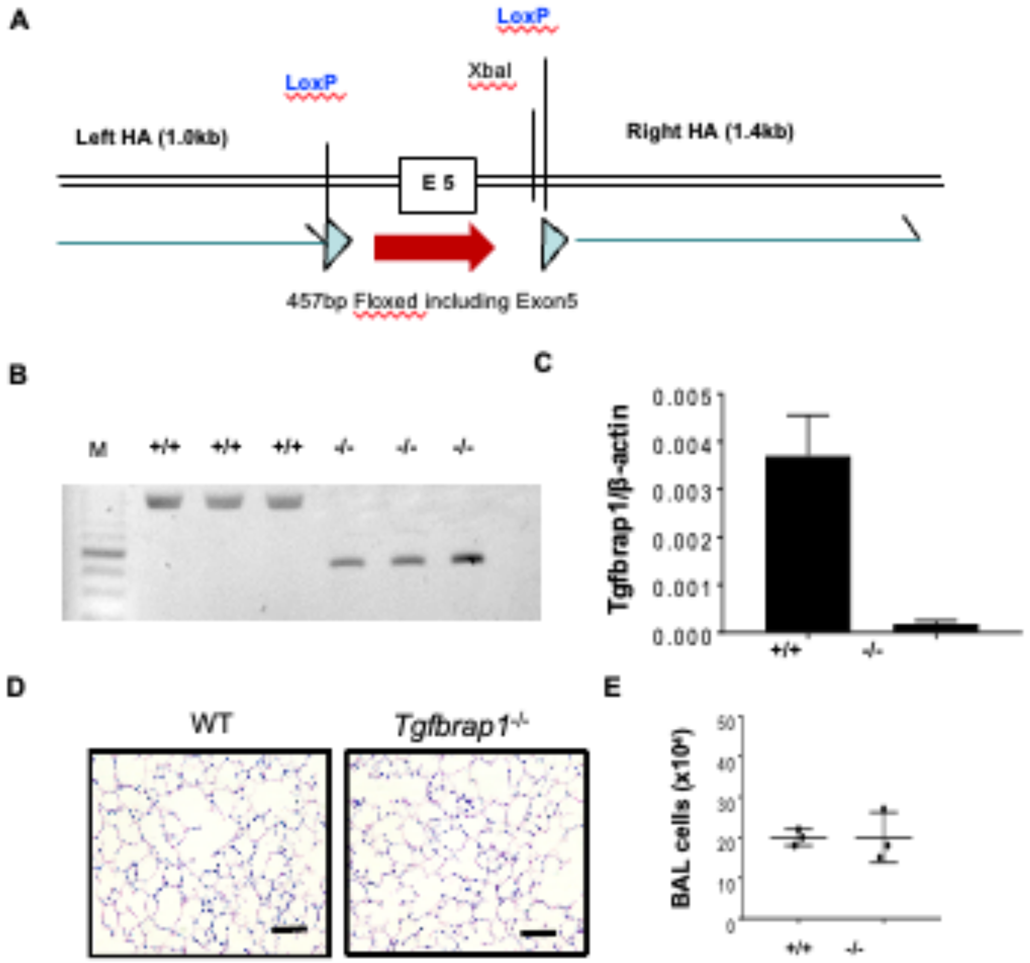


**Figure E2. Generation and basic characterization of Tgfbrap1 KO mice**. (A). Targeting construct, (B) Genotypes of Tgfbrap1 WT and KO mice: +/+, WT; -/-, KO deletion homozygote. (C) Real-time PCR on WT (+/+) and KO (-/-) mice. (D) H&E stains of the lungs that demonstrate no destructive changes between WT vs *Tgfbrap1*$^{-/-}$ mice. (E) Total BAL cells. HA, homologous arm., E5, Exon5. XbaI, restriction site of XbaI, M, 100bp ladder marker. Bar, 50mM. ns, not significant.

**Figure E3**

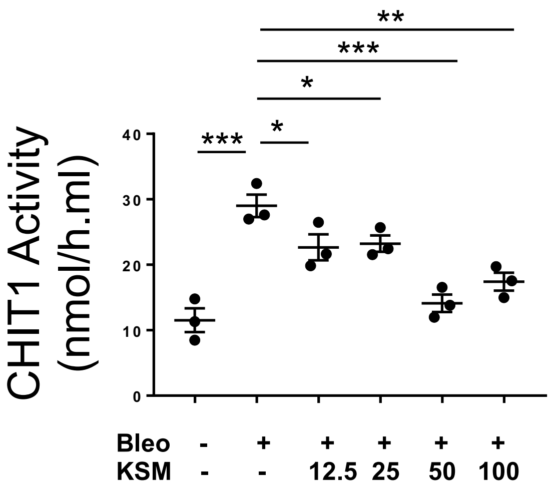


**Figure E3. Anti-CHIT1 activity of KSM in the lungs of bleomycin-challenged mice after KSM administration**. 8 weeks old male mice were challenged with bleomycin (Bleo; i.p, 0.25U/mouse) together with different doses of Kasugamycin (KSM) (12.5 to 100 mg/kg/mouse). Then the mice were sacrificed 2 days after the last KSM treatment, and the lung lysates were prepared and used to evaluate the CHIT1 activity according to the procedures described in the Materials and Methods. **P*<0.05. ***P*<0.01, ****P*<0.001 (One way ANOVA, multiple comparisons)

**Figure E4. Densitometry evaluations on the Western blot images in this study**. For relative quantification of the band intensity of Western blots densitometry data was generated using NIH ImageJ program. The values are mean±SEM. **P*<0.05, ***P*<0.01, ****P*<0.001, *****P*<0.0001 (One way ANOVA, multiple comparisons).

**Fig. 7D**

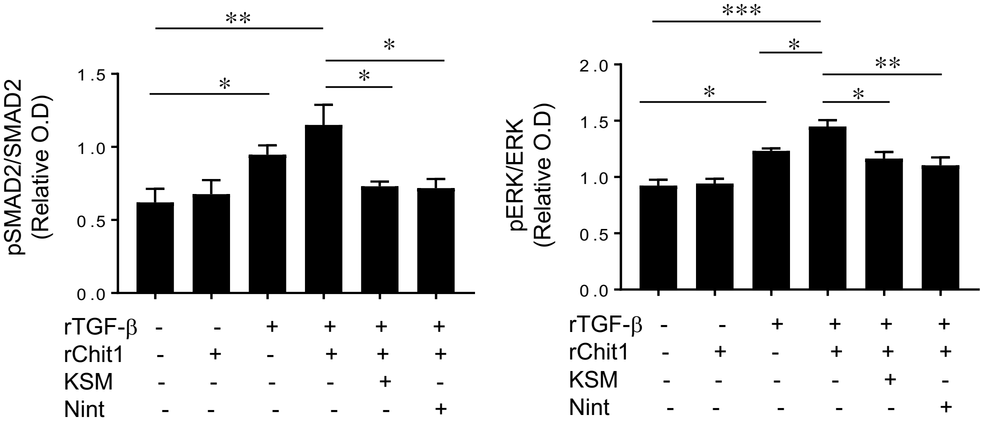


**Fig. 8C**

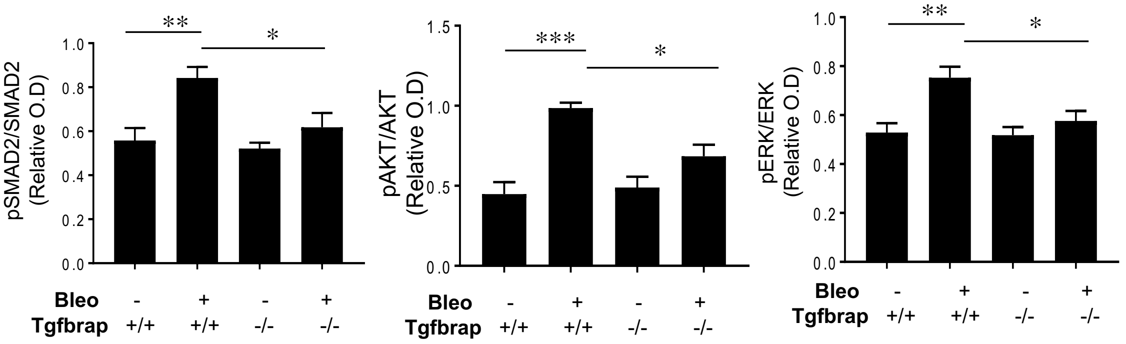


**Fig. 8E**

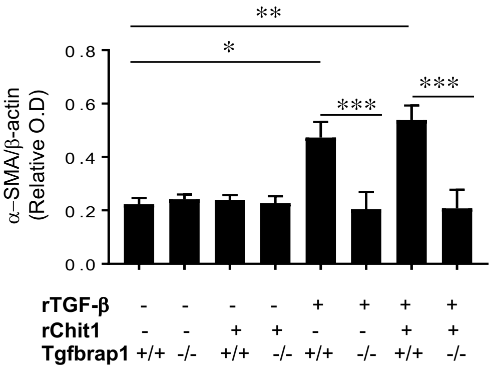

**Figure E4-continued**

**Fig. 8F**

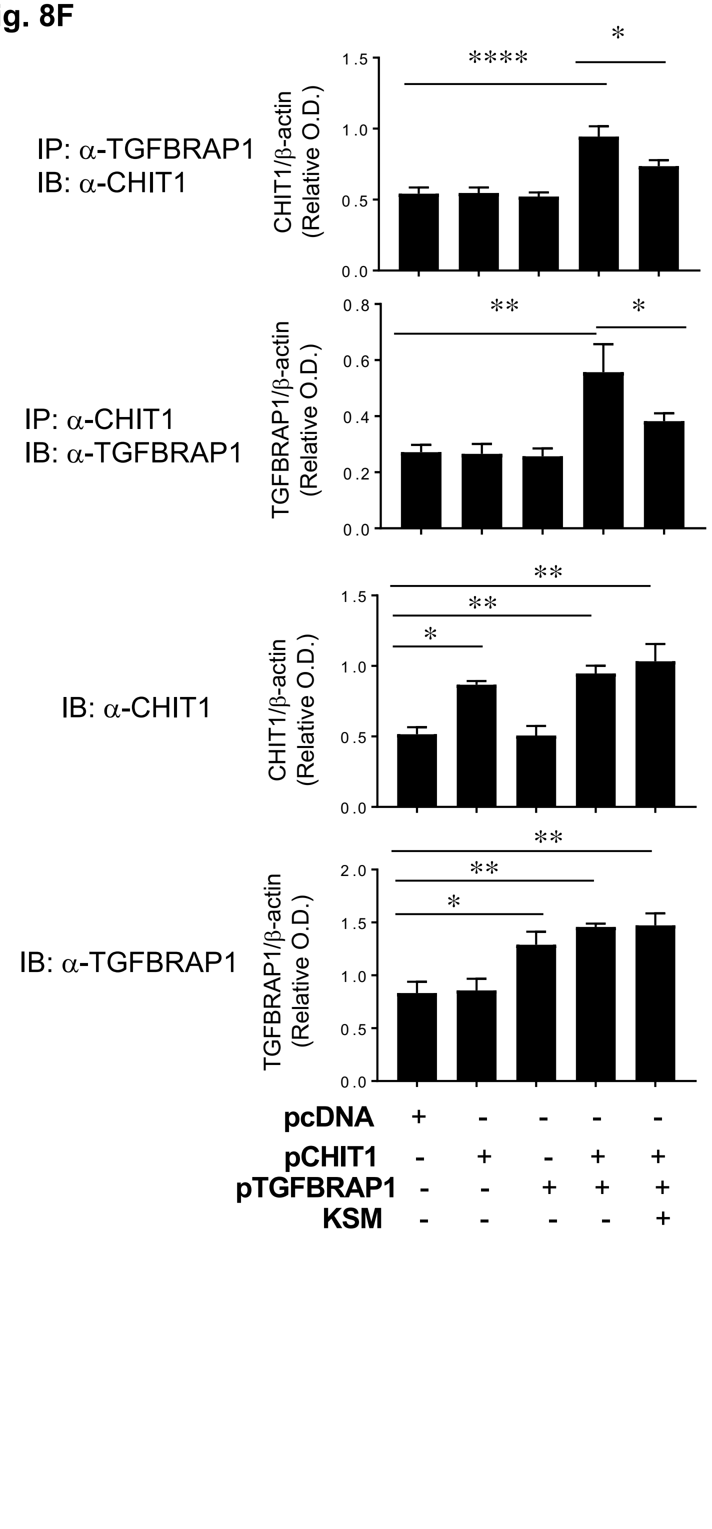
